# Multi-Spatial Scale Dynamic Interactions between Functional Sources Reveal Sex-Specific Changes in Schizophrenia

**DOI:** 10.1101/2021.01.04.425222

**Authors:** A. Iraji, A. Faghiri, Z. Fu, S. Rachakonda, P. Kochunov, A. Belger, J.M. Ford, S. McEwen, D.H. Mathalon, B.A. Mueller, G.D. Pearlson, S.G. Potkin, A. Preda, J.A. Turner, T.G.M. van Erp, V.D. Calhoun

## Abstract

We introduce an extension of independent component analysis (ICA), called multiscale ICA (msICA), and design an approach to capture dynamic functional source interactions within and between multiple spatial scales. msICA estimates functional sources at multiple spatial scales without imposing direct constraints on the size of functional sources, overcomes the limitation of using fixed anatomical locations, and eliminates the need for model-order selection in ICA analysis. We leveraged this approach to study sex-specific and -common connectivity patterns in schizophrenia.

Results show dynamic reconfiguration and interaction within and between multi-spatial scales. Sex-specific differences occur (1) within the subcortical domain, (2) between the somatomotor and cerebellum domains, and (3) between the temporal domain and several others, including the subcortical, visual, and default mode domains. Most of the sex-specific differences belong to between-spatial scale functional interactions and are associated with a dynamic state with strong functional interactions between the visual, somatomotor, and temporal domains and their anticorrelation patterns with the rest of the brain. We observed significant correlations between multi-spatial scale functional interactions and symptom scores, highlighting the importance of multiscale analyses to identify potential biomarkers for schizophrenia. As such, we recommend such analyses as an important option for future functional connectivity studies.

## 1. INTRODUCTION

### 1.1. Multi-Spatial Scale Dynamic Interactions

Brain function has been modeled as coordination and interaction between functional sources, which has been summarized via the principles of segregation and integration (Genon *et al*., 2018). In other words, the brain can be segregated into distinct functional sources (e.g., intrinsic connectivity networks, ICNs), which dynamically interact with each other (i.e., functional integration). Notably, functional sources exist at different spatial scales, and dynamic functional interactions occur both within and between different spatial scales. Previous work has highlighted the importance of analysis at multiple spatial scales (Li *et al*., 2018b); however, most multiple-spatial scale studies have built upon a single set of nodes (e.g., predefined regions or single model order ICA) and identifying multiple levels of modularity (e.g., with different resolution parameter) or clusters (e.g., different number of clusters) (Doucet *et al*., 2011). In the case of using functional sources as nodes, information at different spatial scales captures functional integration among those sources at multiple resolutions. However, each spatial scale also contains its own functional sources with unique functional information. For instance, larger functional sources are not a simple union of smaller functional sources (Figure 1). In addition, functional interactions occur among functional sources across (within and between) different spatial scales (e.g., large networks interact with small networks), which convey important information about the brain as shown in this study. This relationship is effectively ignored if using a single spatial scale to analyze the data.

**Figure 1.**
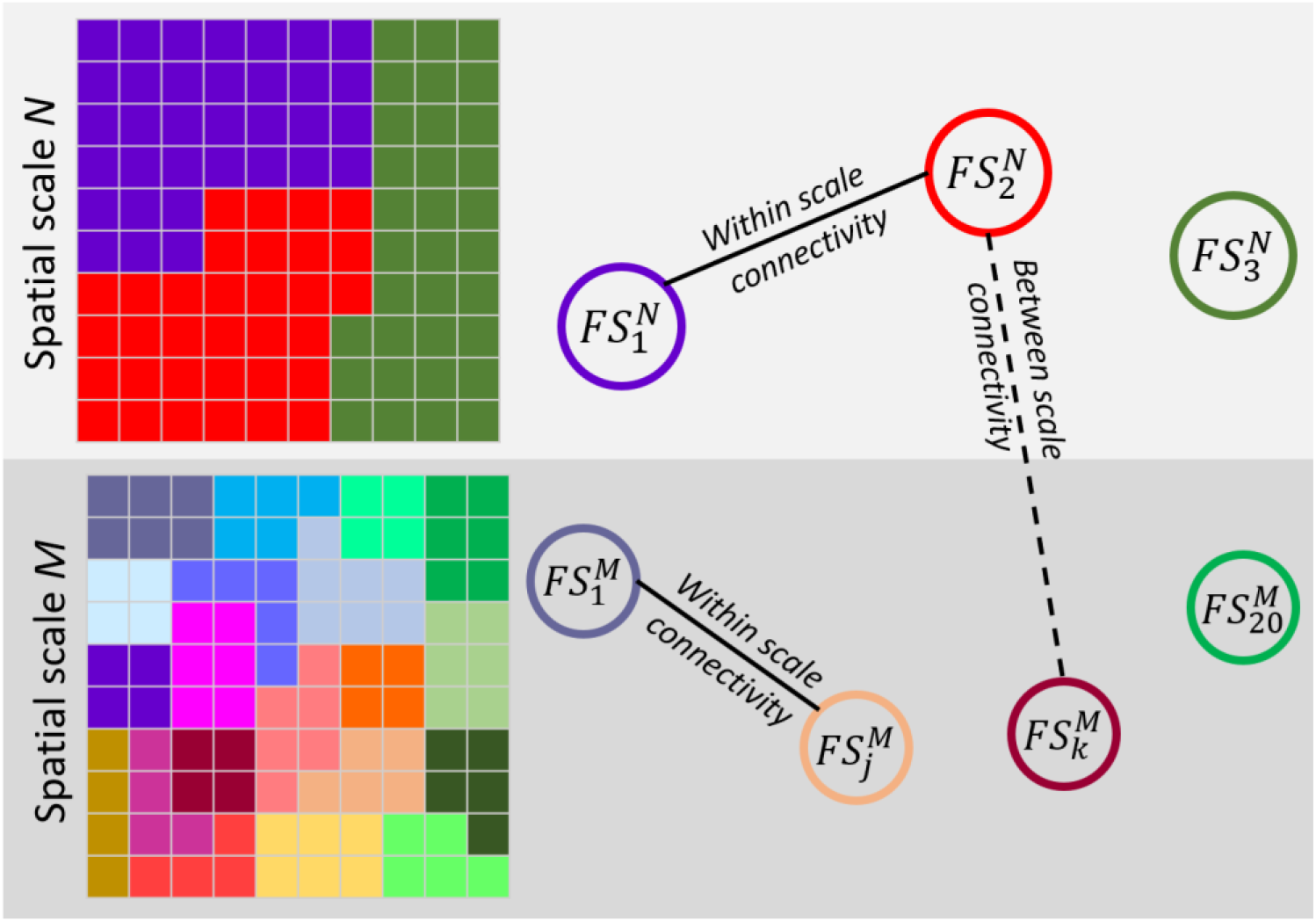
A toy example of multi-spatial scale analysis. A system with two true spatial scales, *M* and *N*. Each spatial scale has its own set of functional source (FSs). Functional sources of spatial scale *M* are not just a simple split of functional sources of scale *N*. Each functional source represents a segregation unit in a given scale, and functional connectivity between functional sources indicates functional integration. Functional interaction (functional connectivity) exist both within and between different spatial scales

Here, we present an approach that combines multiscale ICA (msICA) and functional network connectivity (FNC) to study multi-spatial scale functional interactions (both within and between spatial scales). msICA uses multi-model-order ICA to estimate functional sources at multiple spatial scales directly. Static and dynamic FNC (sFNC/dFNC) were applied to capture static and dynamic interactions between functional sources, both within and between multiple spatial scales. We leveraged this approach to study sex-specific and sex-common schizophrenia differences, which have been understudied but may play an important role in understanding the neural mechanisms as it is clear there are sex differences in schizophrenia, for example in disease onset (Nawka *et al*., 2013; Li *et al*., 2016).

### 1.2. Intrinsic Connectivity Networks (ICNs): Assessment of Functional Sources

A functional source can be defined as a temporally synchronized pattern (Iraji *et al*., 2020b), and studying brain function requires a proper estimation of functional sources to prevent incorrect functional connectivity inferences (Iraji *et al*., 2020b). Due to its emphasis on capturing spatially distinct and temporally coherent sources, ICA has proven itself as a strong method to identify functional source estimates. ICA is a data-driven multivariate technique, which divides the brain into overlapping functionally distinct patterns (Calhoun and Adalı, 2012; Calhoun and de Lacy, 2017; Kaboodvand *et al*., 2018), called intrinsic connectivity networks (ICNs). Each ICN is a temporally synchronized pattern of the brain, a good estimation of a functional source. The ICN time course describes its functional activity over time, while its spatial pattern indicates the contribution of spatial locations to ICN. The spatial scale of ICNs can be set effectively using the model orders of ICA. In other words, we can study brain segregation and estimate ICNs at different spatial scales by using ICA with different model orders. Low-model order ICAs result in large-scale spatially distributed ICNs (Damoiseaux *et al*., 2008; Iraji *et al*., 2016), while higher model order results in more spatially granular ICNs (Allen *et al*., 2011; Iraji *et al*., 2019c; Iraji *et al*., 2019d). Therefore, we proposed to use msICA (running ICA with multiple orders) to estimate functional sources of multiple spatial scales. While there have been a few studies of the effect of model-order on the spatial maps of ICNs (Abou-Elseoud *et al*., 2010), to our knowledge, there is no work which studied brain function across multiple model orders. Similarly, no work has yet evaluated dynamic functional interaction jointly at multiple model orders.

It is worth mentioning that ICA does not impose a direct constraint on the spatial extent of functional sources estimates; thus, msICA allows data itself determined the spatial extend of estimated functional sources without generating spurious sources for different spatial scales. In other words, msICA does not force the functional sources of a given spatial scale to have a similar spatial extent. This gives msICA a great advantage as we do not expect different brain areas or functional domains, e.g., “the primary cortex vs. the frontal lobe” and “the visual domain vs. the cognitive control domain across” to be parceled at the same level of granularity (see Section 3.1).

msICA also addresses the model-order selection problem as, in general, one remedy to parameter selections is finding a procedure to combine results from several parameters. Various information-theoretic criteria such as the Minimum description-length criterion (MDL) and Akaike’s information criterion (AIC) have been used to estimate an optimal model order. However, the optimal number can vary across them; as such, the model-order selection problem still remains as selecting estimation method. Instead of focusing on a single model order selected by these approaches, msICA includes information of all spatial scales (within the constraints of the number of model orders we use).

### 1.3. Functional Network Connectivity (FNC): Assessment of Functional Interaction

While ICA effectively segregates the brain into ICNs, FNC provides a way to study functional interaction and integration. FNC is defined as the temporal dependency among ICNs and commonly estimated using Pearson’s correlation coefficient between ICN time courses (Jafri *et al*., 2008). Thus, FNC characterizes the functionally integrated relationship across the brain by calculating the functional interaction between ICNs.

Traditionally, functional integration has been studied using sFNC, where the overall functional interactions are calculated using scan-length averaged FNC. However, the brain constantly integrates and processes the information in real-time. Considering the brain’s rich, dynamic nature, a number of methods have moved beyond the “static” oversimplification and evaluate the temporal reconfiguration of functional interactions using dFNC (Allen *et al*., 2014; Calhoun *et al*., 2014; Iraji *et al*., 2020a). The dFNC approaches calculate time-resolved FNC allowing us to study variations in functional integrations over time and identify different brain functional interaction patterns, also known as brain functional states (Iraji *et al*., 2020a).

### 1.4. Schizophrenia

Schizophrenia is a psychotic disorder accompanied by various cognitive impairments and a decrease in social and occupational functioning. Schizophrenia is a heterogeneous syndromic diagnosis of exclusion, lacks unique symptoms and is diagnosed clinically by both positive symptoms, such as delusions, hallucinations, disorganized speech, disorganized or catatonic behavior; and negative symptoms, such as apathy, blunted affect, and anhedonia (Association, 2013), plus a decline in social functioning. Schizophrenia overlaps considerably with both schizo-affective disorder and psychotic bipolar disorder, not only symptomatically, but in terms of genetics and at the level of other biomarkers (Clementz *et al*., 2016). The diverse temporal trajectory across individuals with SZ and the different types of clinical symptoms suggest alterations in various functional domains and brain capacity reductions to integrate information across the brain. Schizophrenia has been hypothesized as a developmental disorder of disrupted brain function, which can be characterized by functional dysconnectivity and/or changes in functional integration (Friston and Frith, 1995; Stephan *et al*., 2006; Kahn *et al*., 2015). Therefore, studying static and dynamic FNC can provide vital information about brain functional integration and its schizophrenia changes, potentially improving our understanding of the actual brain pathology underlying different schizophrenia subcategories.

In early work, Meda et al. show abnormal FNC, including those related to paralimbic circuits, which were correlated significantly with PANSS negative scores (Meda *et al*., 2012). Focusing on the default mode, hypoconnectivity was observed across all related networks (Meda *et al*., 2014). Dynamic studies also identify hypoconnectivity as the dominant dysconnectivity pattern, while identifying few consistent hyperconnectivity patterns. The strengths of dFC between subcortical and sensory networks are weaker in individuals with schizophrenia (Damaraju *et al*., 2014). The weaker dynamic functional connectivity (dFC) strengths have also been observed in several brain networks in spatial dynamic studies (Iraji *et al*., 2019a; Iraji *et al*., 2019d). The decrease in the strengths of dFC (transient hypoconnectivity) seems to be accompanied by higher fluctuations of dFC between brain regions (Yue *et al*., 2018) and within and between several brain networks (Ma *et al*., 2014; Iraji *et al*., 2019a). Sun et al. reported overall higher global efficiency across the schizophrenia brain (Sun *et al*., 2019). The alteration in the dFNC patterns in schizophrenia also seems to be related to cognitive performance (Fu *et al*., 2018; Yue *et al*., 2018; Iraji *et al*., 2019a). For instance, the temporal variability of FNC between the amygdala-medial prefrontal cortex (mPFC) is positively correlated with total symptom severity and negatively correlated with information processing efficiency (Yue *et al*., 2018). The correlation between the energy index (spatiotemporal uniformity) of the subcortical domain and the attention/vigilance domain of computerized multiphasic interactive neurocognitive dualdisplay system (CMINDS) was reported to be disrupted in schizophrenia (Iraji *et al*., 2019a). Studies also show frequency-specific dFC alterations in SZ patients (Yaesoubi *et al*., 2017; Zhang *et al*., 2018; Faghiri *et al*., 2020b). However, previous studies have not studied functional interactions across multiple spatial scales and have underappreciated differences between male and female cohorts (Damaraju *et al*., 2014; Miller *et al*., 2016; Iraji *et al*., 2019a; Miller *et al*., 2019; Faghiri *et al*., 2020b).

Schizophrenia incidence is higher in men (Aleman *et al*., 2003; McGrath *et al*., 2004), but paradoxically there is equal overall prevalence (Saha *et al*., 2005). There is also evidence suggesting sex-differences in onset, symptom expression, and outcome in schizophrenia (Navarro *et al*., 1996; Nawka *et al*., 2013; Li *et al*., 2016). For instance, males have more severe overall symptoms, worse outcomes, more negative and fewer affective symptoms, and experience symptoms earlier than females (Li *et al*., 2016). Furthermore, symptoms respond more quickly to treatments in females. However, sex differences in symptoms and outcomes also depend on the age of onset and treatment (Li *et al*., 2016; Seeman, 2019). Understanding sex-specific characteristics of functional connectivity, which is currently lacking in the field, can help provide an important insight to understand sex differences in schizophrenia and potentially the opportunity to deliver sex-specific treatments and care for individuals with schizophrenia.

Considering the previous static and dynamic FNC findings on sex differences in typical control cohorts (Allen *et al*., 2011; Yaesoubi *et al*., 2020) and previous report on sex differences in schizophrenia (Navarro *et al*., 1996; Nawka *et al*., 2013; Li *et al*., 2016), we hypothesize that multiscale functional interactions capture sex-specific changes in schizophrenia, which are significantly correlated with schizophrenia’s symptoms score. We examined our hypothesis using the following pipeline: 1) we estimated ICNs at multiple spatial scales using ICA with model-orders of 25, 50, 75, and 100; 2) we calculated the multi-spatial scale static and dynamic functional integrations using within and between model-orders sFNC and dFNC using a window-based approach (Allen *et al*., 2014; Iraji *et al*., 2020a); 3) we evaluated sex-specific differences between typical controls and individuals with schizophrenia.

## 2. MATERIALS AND METHODS

### 2.1. Participant Demographics and Data Selection Inclusion Criteria

The data used in this study selected from three projects, including FBIRN (Functional Imaging Biomedical Informatics Research Network), MPRC (Maryland Psychiatric Research Center), and COBRE (Center for Biomedical Research Excellence). We selected a subset of data that satisfies the inclusion criteria, including 1) data of individual with typical control or schizophrenia diagnosis; 2) data with high-quality registration to echo-planar imaging (EPI) template; and 3) the head motion transition should be less than 3° rotations and 3 mm translations in every direction (Fu *et al*., 2020). Mean framewise displacement among selected subject is average ± standard deviation = 0.1778 ± 0.1228; min ~ man = 0.0355 ~ 0.9441. Thus, we report on resting-state fMRI (rsfMRI) data from 827 individuals, including 477 typical controls and 350 individuals with schizophrenia selected (Table 1).

**Table 1.** Demographic information of the data used in the study. FBIRN: Functional Imaging Biomedical Informatics Research Network, MPRC: Maryland Psychiatric Research Center, COBRE: Center for Biomedical Research Excellence

| Project | diagnostic | N | sex | N | Age (years) |  |
| --- | --- | --- | --- | --- | --- | --- |
| | | | | | mean ± sd | median/range |
| FBIRN | Control group | 160 | Male | 115 | 37.26 ± 10.71 | 39/(19-59) |
| | | | Female | 45 | 36.47 ± 11.33 | 33/(19-58) |
| | Schizophrenia group | 150 | Male | 114 | $38.74 \pm 11.78$ | 40/(18-62) |
| | | | Female | 36 | $39.06 \pm 11.40$ | 36/(21-57) |
| **MPRC** | Control group | 238 | Male | 94 | $38.72 \pm 13.63$ | 40/(12-68) |
| | | | Female | 144 | $41.22 \pm 16.06$ | 44/(10-79) |
| | Schizophrenia group | 150 | Male | 98 | $35.57 \pm 13.18$ | 32/(13-63) |
| | | | Female | 52 | $44.60 \pm 13.87$ | 47/(13-63) |
| **COBRE** | Control group | 79 | Male | 55 | $39.07 \pm 12.43$ | 38/(18-65) |
| | | | Female | 24 | $34.92 \pm 10.23$ | 34/(18-58) |
| | Schizophrenia group | 50 | Male | 42 | $37.43 \pm 15.05$ | 32.5/(19-64) |
| | | | Female | 8 | $43.25 \pm 12.78$ | 40/(31-65) |

### 2.2. Data Acquisition

The FBIRN dataset was collected from seven sites. The same resting-state fMRI (rsfMRI) parameters were used across all sites: a standard gradient echo-planar imaging (EPI) sequence, repetition time (TR)/echo time (TE) = 2000/30 ms, voxel spacing size = 3.4375 × 3.4375 × 4 mm, slice gap = 1 mm, flip angle (FA) = 77°, field of view (FOV) = 220 × 220 mm, and a total of 162 volume. Six of the seven sites used 3-Tesla Siemens Tim Trio scanners, and one site used a 3-Tesla General Electric Discovery MR750 scanner.

The MPRC dataset was collected in three sites using a standard EPI sequence, including Siemens 3-Tesla Siemens Allegra scanner (TR/TE = 2000/27 ms, voxel spacing size = 3.44 × 3.44 × 4 mm, FOV = 220 × 220 mm, and 150 volumes), 3-Tesla Siemens Trio scanner (TR/TE = 2210/30 ms, voxel spacing size = 3.44 × 3.44 × 4 mm, FOV = 220 × 220 mm, and 140 volumes), and 3-Tesla Siemens Tim Trio scanner (TR/TE = 2000/30 ms, voxel spacing size = 1.72 × 1.72 × 4 mm, FOV = 220 × 220 mm, and 444 volumes).

The COBRE dataset was collected in one site using a standard EPI sequence with TR/TE = 2000/29 ms, voxel spacing size = 3.75 × 3.75 × 4.5 mm, slice gap = 1.05 mm, FA = 75°, FOV = 240 × 240 mm, and a total of 149 volumes. Data was collected using a 3-Tesla Siemens TIM Trio scanner.

### 2.3. Preprocessing/MRI Data Preprocessing

The preprocessing was performed primarily using the statistical parametric mapping (SPM12, http://www.fil.ion.ucl.ac.uk/spm/) toolbox. The rsfMRI data preprocessing using the following steps: 1) discarding the first five volumes for magnetization equilibrium purposes, 2) rigid motion correction to correct subject head motion during scan, and 3) slice-time correction to account for temporally misalignment in data acquisition. Next, the data of each subject was nonlinearly registered to a Montreal Neurological Institute (MNI) echo-planar imaging (EPI) template, resampled to 3 mm^3^ isotropic voxels, and spatially smoothed using a Gaussian kernel with a 6 mm full width at half-maximum (FWHM = 6 mm). The voxel time courses were then z-scored (variance normalized). We are interested in identifying functional sources, temporally synchronized regions. Therefore, temporal coupling and not amplitude information are the information of interest. Variance normalization was shown to enhance sensitivity to functional segregation and functional sources (Iraji *et al*., 2019d) and be highly reproducible across different studies. Furthermore, prior to calculating static and dynamic FNC, an additional post hoc cleaning procedure was performed on the time courses of ICNs to reduce the effect of remaining noise, which may not be wholly removed using ICA, and to improve the detection of dynamic FNC patterns (Allen *et al*., 2014). ICNs time courses were detrended by removing linear, quadratic, and cubic trends. The six motion realignment parameters and their derivatives were regressed out. Outliers were detected based on the median absolute deviation, similar to implemented in AFNI 3Ddespike (http://afni.nimh.nih.gov/), and replaced with the best estimate using a third-order spline fit to the clean portions of the time courses. Bandpass filtering was applied using a fifth-order Butterworth filter with a cutoff frequency of 0.01 Hz-0.15 Hz.

### 2.4. Intrinsic Connectivity Network (ICNs) Estimation

For the initial work in this paper, we utilized spatial ICA with several model orders (25, 50, 75, and 100) to identify intrinsic connectivity networks (ICNs) at multiple spatial scales. Similar to most ICA-based studies of fMRI, we implemented group-level spatial ICA followed by a back-reconstruction technique to estimate subjects-specific independent components (ICs) time courses.

We used the GIFT toolbox (https://trendscenter.org/software/gift/) (Calhoun *et al*., 2001; Calhoun and Adali, 2012; Iraji *et al*., 2020a). First, subject-specific spatial principal components analysis (PCA) was applied to normalize the data and to allow subjects to contribute similarly in the common subspace. The subject-specific PCA also has denoising and computational benefits (Erhardt *et al*., 2011). We retain maximum subject-level variance (greater than 99.99%). While the subject-specific PCA privileges subject differences at the subject-level, the group-level PCA favors subject commonalities (Erhardt *et al*., 2011). All subject-level principal components were concatenated together across the time dimension, and group-level spatial PCA was applied to concatenated subject-level principal components. *N* (25, 50, 75, and 100) group-level principal components that explained the maximum variance were selected as the input for spatial ICA to calculate *N* (25, 50, 75, and 100) group independent components.

Infomax was chosen as the ICA algorithm because it has been widely used and compares favorability with other algorithms (Correa et al., 2007a; Correa et al., 2005). For each model-order (N = 25, 50, 75, and 100), the Infomax ICA algorithm was run 100 times and clustered together within the ICASSO framework (Himberg *et al*., 2004). The run with the closest independent components to the centrotypes of stable clusters (ICASSO cluster quality index > 0.8) was selected as the best run and used for future analysis (Ma *et al*., 2011b). This is an important point and facilitates replicable results. Next, the subject-specific ICs time courses were calculated using the spatial multiple regression technique (Calhoun *et al*., 2004). At each time point, the contribution of each IC to the BOLD signal was calculated using linear regression (Calhoun *et al*., 2004).

We selected a subset of independent components as ICNs if they are stable (ICASSO stability index > 0.8) and depict common ICNs properties including 1) dominant low-frequency fluctuations of their time courses evaluated using dynamic range and the ratio of low frequency to high-frequency power, 2) exhibit peak activations in the gray matter, 3) have low spatial overlap with vascular, ventricular, and 4) low spatial similarity with motion and other known artifacts. Finally, ICNs were grouped into functional domains based on prior knowledge of their anatomical and functional properties (Allen, Erhardt et al. 2011).

### 2.5. Static and Dynamic FNC Calculation

We calculated static and dynamic functional network connectivity (FNC) between every single pair of ICNs across all model-orders to effectively capture functional integration and interaction across different spatial scales. For a subset of data (15%) with a sampling rate different than 2 seconds, ICNs time courses were interpolated to 2 seconds. Minimum data length across all subjects was selected for further analysis. Static FNC (sFNC) was estimated by calculating the Pearson correlation between each pair of ICNs time courses resulting in one sFNC matrix for each individual. Each element of the sFNC matrix is the functional connectivity between a pair of ICNs.

In contrast to sFNC, which uses the full length of scan, in dynamic FNC (dFNC), we calculate multiple FNC matrices for different time segments of scan (i.e., FNC matrices for durations smaller than the whole time series) (Iraji *et al*., 2020a). As a result, we can study variations in FNC over time. Here, we used a windowed-based approach with the slide step size of two seconds (maximum overlap between consecutive windows). A recommended window size is between 30 and 60 seconds (Iraji *et al*., 2020a); thus, we chose the middle value (44 seconds, timepoint increment is 2 seconds). Tapered window created by convolving a rectangle window (width = 44 seconds) with a Gaussian (σ = 6 seconds) and used to calculate windowed-FNC. This results in a series of windowed-FNC matrices over time (FNC as a function of time) containing dFNC information.

Next, we identified dFNC states from windowed-FNC matrices using the k-means clustering, in which each cluster represents one dynamic state (Iraji *et al*., 2020a). We applied a two-stage k-means clustering. First, windows with local maxima with FNC variances were selected for each subject, and k-means clustering was applied to the set of all subject-specific local maxima (also known as exemplars). We used the city-block distance metric because it is suggested to be a more effective dissimilarity measure than Euclidean distance for high-dimensional data (Aggarwal *et al*., 2001). K-means clustering was repeated 100 times with different initializations using the k-means++ technique to increase the chance of escaping local minima. The resulting centroids were then used to initialize a clustering to all 93,451 (827 subjects × 113 windows) windowed-FNC matrices. The optimal number of dFNC states was selected based on the elbow criterion by calculating the ratio of within to between cluster variance and running the clustering procedure for 1 to 15 clusters. Subject-specific dFNC states were next estimated by averaging windowed-FNC of time-windows assigned to a given state.

We repeated the dFNC states identification procedures using two alternative ways to ensure that the dFNC states are not biased to the clustering algorithm. 1) We first applied k-means clustering at the subject-level and then concatenated the subject-level centroids for group-level clustering and identifying dFNC states. 2) We directly applied k-mean clustering to all 93,451 (827 subjects × 113 windows) windowed-FNC matrices. We also evaluated the clustering results using Euclidean and Correlation distances.

### 2.6. Group Comparison Analysis

We evaluated sex-specific differences in multiscale sFNC and dFNC between the control group (CT) and the individuals with schizophrenia (SZ). For each sex cohort, male and female, we separately assessed diagnostic group differences, i.e., male controls versus male individuals with schizophrenia (maleCT vs. maleSZ) and female controls versus female individuals with schizophrenia (femaleCT vs. femaleSZ). We used a general linear model (GLM) with age, data acquisition site, and mean framewise displacement as covariates. Framewise displacement is the sum of changes in the six rigid-body transform parameters (framewise displacement(t) = |Δdx(t)| + |Δdy(t)| + |Δdz(t)| + |Δα(t)| + |Δβ(t)| + |Δγ(t)|). Mean framewise displacement was added to the GLM to account for any residual motion effect that was not removed in the previous three motion-removal steps. The statistical analysis results were corrected for multiple comparisons using a 5% false discovery rate (FDR). It is worth mentioning that all statistical analysis results were combined (sFNC and dFNC; male and female; across all model orders) and corrected for multiple comparisons, which is more conservative than correcting for each statistical analysis separately.

Next, we evaluated sex-specific differences for the sFNC and dFNC features that showed a significant difference between the control group and individuals with schizophrenia in either of the sex cohorts (“maleCT vs. maleSZ” and/or “femaleCT vs. femalesSZ”). For each feature, we compared the difference of the *t*-value of GLM statistic between two sex cohorts (“*t*-value of maleCT vs. maleSZ” - “*t*-value of femaleSZ vs. femaleCT”) with a null distribution. The *p*-value of the *t*-value of difference was corrected for multiple comparisons using the same procedure explained in the previous paragraph.

The null distribution was created by randomly permuting sex labels within each diagnostic group. In other words, the diagnostic label remained intact; individuals with schizophrenia remained schizophrenia, and control subjects remained in the control group, and only the sex labels were randomly permuted. Furthermore, the number of females and males in each diagnostic group did not change. This permutation process was repeated 5000 times. For each permutation, the GLM was applied to two null male and null female cohorts independently. For each feature, the difference of the *t*-value of diagnosis for two null cohorts was calculated. This results in 5000 samples of the null distribution for each feature.

We also studied sex-specific differences at the domain-level across different spatial scales. For static FNC and each dynamic state, the average FNC was calculated within and between seven functional domains both within and between four model-orders (e.g., “CC25-DM25 and “CR50-VS100”). For example, “CR50-VS100” is the average FNC between every pair of ICNs belong to the cerebellum domain model order 50 and ICNs belong to the visual domain, model order 100. This results in a 28-by-28 domain-level functional integration matrix. The static and dynamic state domain-level functional integration matrices were then evaluated for sex-specific differences.

### 2.7. Relationship with Symptom Scores

We further evaluate if the multiscale functional network connectivity pairs showing sex-specific changes in schizophrenia are related to the symptoms of schizophrenia. The positive and negative syndrome scale (PANSS) scores are available for the FBIRN dataset, while the MPRC dataset includes the brief psychiatric rating scale (BPRS) scores. We transformed BPRS total scores to PANSS total scores using the matching obtained from 3767 individuals (Leucht *et al*., 2013). Next, we evaluated the relationship between the PANSS total score and domain-level features with significant sex-specific differences. Correlation analyses were conducted after regressing out age, site, and meanFD and corrected for multiple comparisons.

## 3. RESULTS

### 3.1. Multi-Spatial Scale Functional Segregation: Intrinsic Connectivity Networks (ICNs)

We performed spatial ICA with 25, 50, 75, and 100 components on rsfMRI data from 827 subjects to functionally segregate the brain at different spatial scales. Based on the criteria explained in Section 2.4, we identified 15, 28, 36, and 48 independent components as ICNs for model orders 25, 50, 75, and 100, respectively. Detailed information of the ICNs, including spatial maps, coordinates of peak activations, and temporal and frequency information, can be found in Supplementary 1. ICNs were grouped into seven functional domains (FDs), including Cognitive Control (CC), Cerebellum (CR), Default Mode (DM), Subcortical (SB), Somatomotor (SM), Temporal (TP), Visual (VS). Figure 2 illustrates the composite views of functional domains for each model order and aggregated. Each composite view is obtained by thresholding and overlaying associated ICNs. For example, the first image in subplot(CR,ICA25) was obtained by thresholding (|Z|>1.96) and overlaying two ICNs associated with the cerebellum domain in model order 25. Table 2 shows the number of ICNs for each model order and functional domain. The results suggest as the model order increases, the number of ICNs increases, and the brain and the functional domains segregate into more functional sources (ICNs). For instance, the subcortical (SB) domain consists of only one ICN in model order 25, enclosing the whole subcortical regions, while it parcels into spatially distinct ICNs as model order increases. However, the number of ICNs does not increase proportionally with model order. While some functional domains break into more ICNs as the model order increases, others demonstrate a smaller amount of changes in the number of ICNs and their spatial distributions across model-orders studied in this work. For example, we observe significant changes in the ICNs associated with the cognitive control domain across model orders, particularly between model order 50 and 75, while the number of ICNs are the same for model order 50 and 75 for the somatomotor and visual domains. Interestingly, across different model orders, we observed ICNs with high spatial overlap (high spatial similarity) but clearly distinct features. The second row of Figure 5 shows two distinct ICNs with high spatial overlap associated with the primary motor cortex.

**Figure 2.**
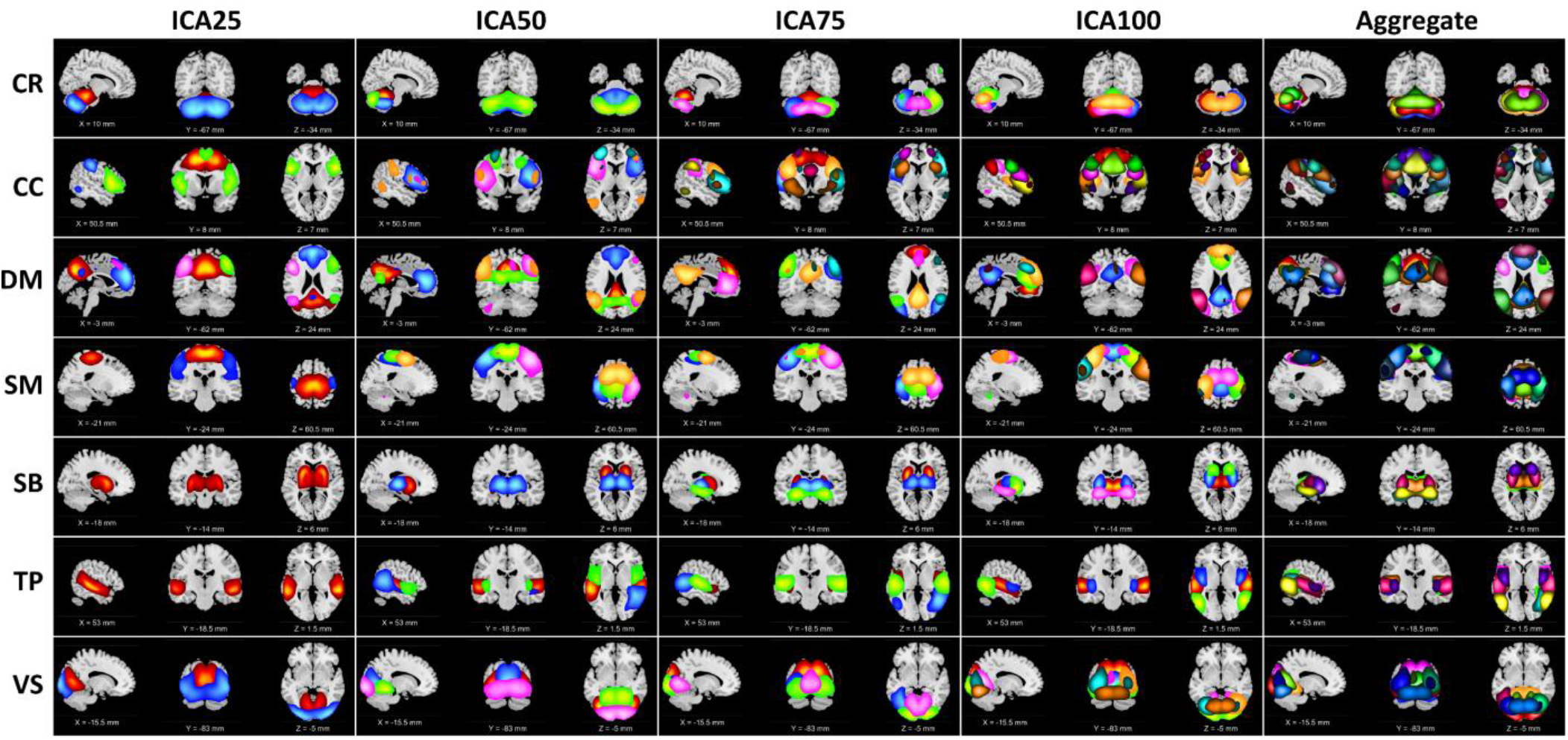
Visualization of the intrinsic connectivity networks (ICNs) identified from four ICA model orders of 25, 50, 75, and 100. ICNs were groups into seven functional domains (FD) based on their anatomical and functional properties. The functional domains are Cognitive Control (CC), Default Mode (DM), Visual (VS), Subcortical (SB), Cerebellum (CR), Somatomotor (SM), and Temporal (TP). Columns represent the composite maps of seven functional domains for four ICA model orders and aggregated. Each color represents the spatial map of one ICN thresholded at |Z| > 1.96 (*p* = 0.05).

**Table 2.**
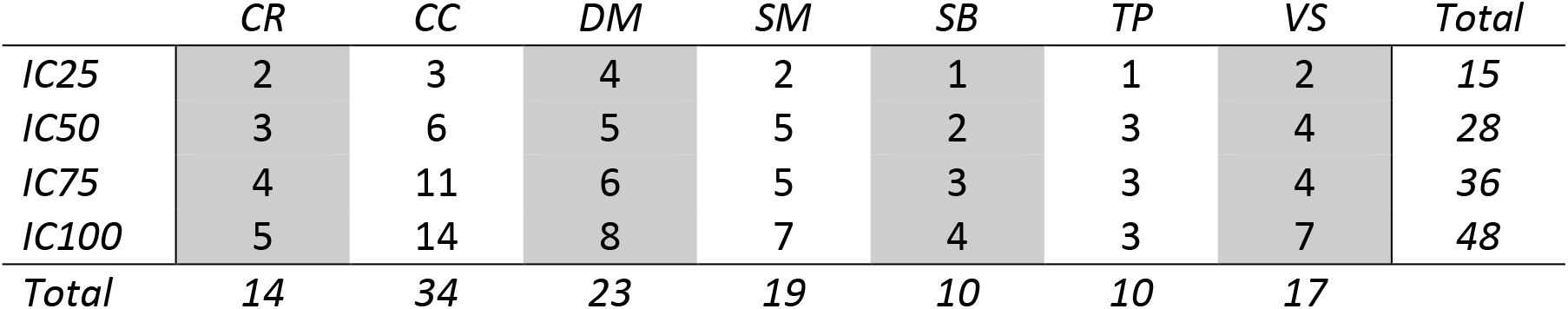
The number of Intrinsic Connectivity Networks (ICNs) for each model order and functional domains, Cognitive Control (CC), Cerebellum (CR), Default Mode (DM), Subcortical (SB), Somatomotor (SM), and Temporal (TP).

### 3.2. Dynamic Functional Integration: Static/Dynamic Functional Network Connectivity (sFNC/dFNC)

*Figure 3(A, I)* and *Figure 4(A, I)* display “block” and “finger” plots of the group-level multiscale functional integration computed using the entire scan length (i.e., static functional network connectivity (sFNC)). sFNC shows similar patterns for control groups, individuals with schizophrenia, males, and females. In the block plot, we sort ICNs by functional domain and then model order. The block plot of sFNC resembles previous single model order studies, showing modular organization within functional domains across model orders. Consistent with prior literature (Allen *et al*., 2011), we observed an overall negative association (anticorrelation) between the default model and the rest of the brain, particularly the visual, somatomotor, and temporal domains, during rest. Interestingly, this negative association with more prominent between model orders, for example, between the default mode of model order 25 (DM25) and the somatomotor of model order 100 (SM100). We also observed strong FNC between the somatomotor, temporal, and visual domains, and between the subcortical and cerebellum domains. Figure 3 suggests that the FNC within functional domains is stronger than between functional domains, and this pattern is consistent for both within and between model orders. The similarity in FNC pattern within and between model orders can be observed in the finger plots (Figure 4), where ICNs are sorted first by model order and then functional domains. The finger plot (Figure 4) shows functional domain (FD) modular patterns (stronger FNC within FDs compared to between FDs) between model orders similar to within model order.

**Figure 3.**
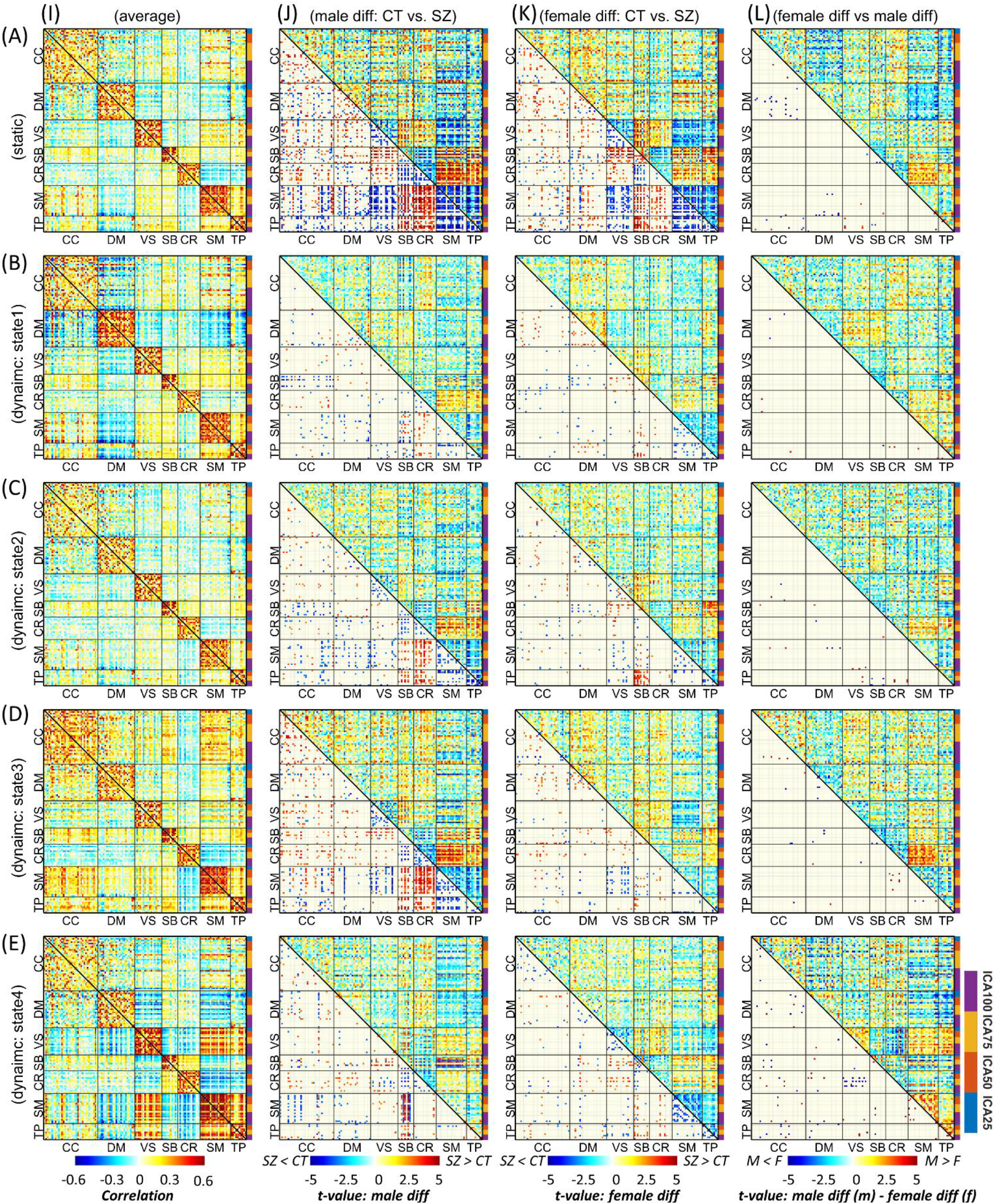
Block plot of multiscale functional integration. ICNs are sorted by functional domain and then model order. Row *A* is the result of static FNC analysis, and rows *B* to *E* represent the four dynamic states. Column *I* is the average FNC matrix for static FNC and dynamic FNC states. Column *J* shows the result of group comparison between male individuals with schizophrenia (SZ) and the male control group (CT). Column *K* shows the result of group comparison between SZ and CT individuals in the female cohort. In columns *J* and *K*, the upper triangular shows the *t*-value of statistical comparisons, and the lower triangular shows statistically significant differences after FDR correction for multiple comparisons (FDR-corrected threshold = 0.05). Column *L* shows the result of the statistical comparison between the differences observed in the male cohort versus the female cohort. The upper triangular in shows the differences between the *t*-value of statistical comparisons in male and female cohorts (“*t*-value of maleSZ vs. maleCT” - “*í*-value of femaleSZ vs. femaleCT”), and the lower triangular shows the SZ-associated abnormal patterns that are significantly different between male and female cohorts after FDR correction. Cognitive Control (CC), Default Mode (DM), Visual (VS), Subcortical (SB), Cerebellum (CR), Somatomotor (SM), and Temporal (TP).

**Figure 4.**
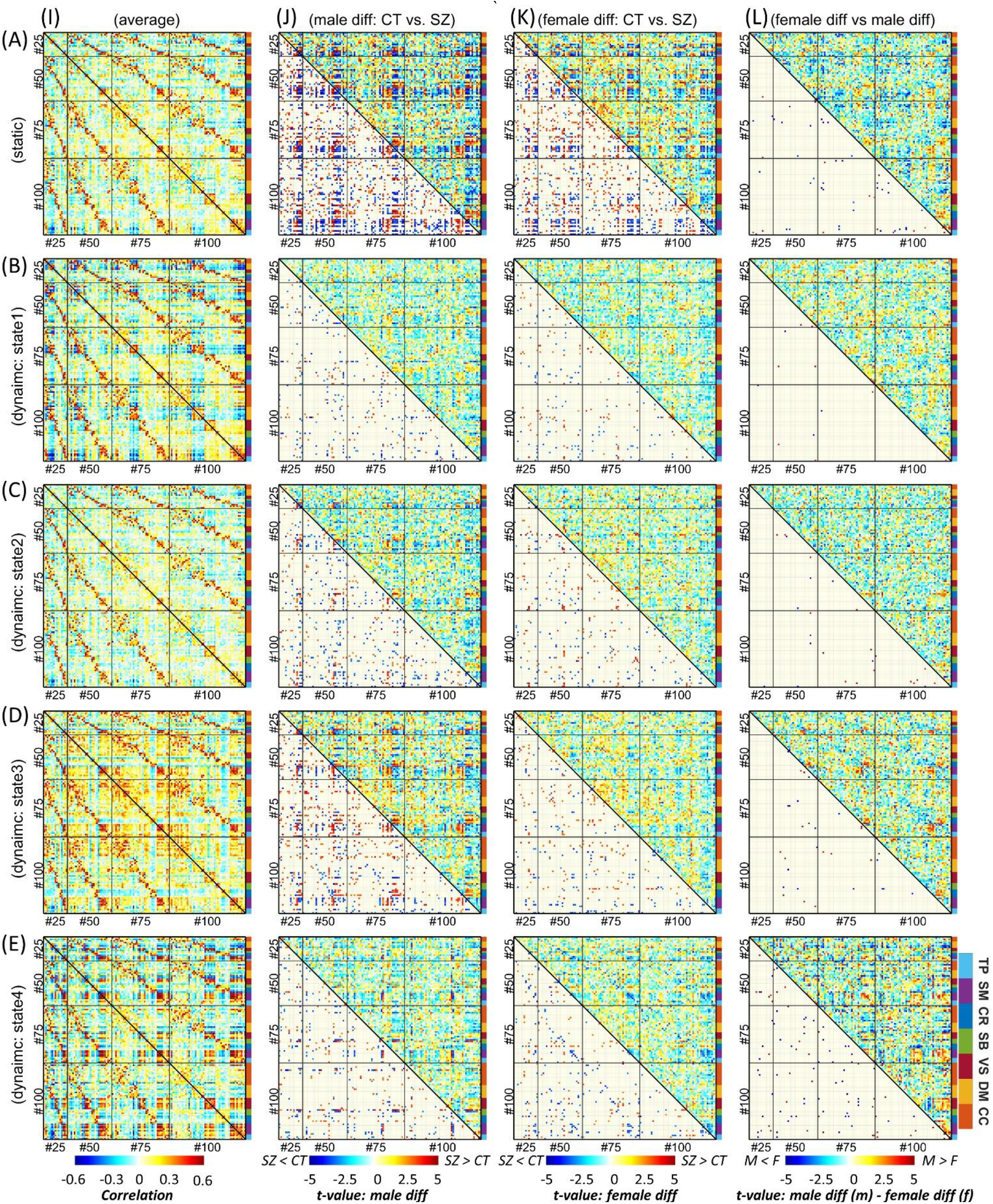
Finger plot of multiscale functional integration. ICNs are sorted by model order then functional domain. Row *A* is the result of static FNC analysis, and rows *B* to *E* represent the four dynamic states. Column *I* is the average FNC matrix for static FNC and dynamic FNC states. Column *J* shows the result of group comparison between male individuals with schizophrenia (SZ) and the male control group (CT). Column *K* shows the result of group comparison between SZ and CT individuals in the female cohort. In columns *J* and *K*, the upper triangular shows the *t*-value of statistical comparisons, and the lower triangular shows statistically significant differences after FDR correction for multiple comparisons (FDR-corrected threshold = 0.05). Column *L* shows the result of the statistical comparison between the differences observed in the male cohort versus the female cohort. The upper triangular in shows the differences between the *t*-value of statistical comparisons in male and female cohorts (“*t*-value of maleSZ vs. maleCT” - “*t*-value of femaleSZ vs. femaleCT”), and the lower triangular shows the SZ-associated abnormal patterns that are significantly different between male and female cohorts after FDR correction. Cognitive Control (CC), Default Mode (DM), Visual (VS), Subcortical (SB), Cerebellum (CR), Somatomotor (SM), and Temporal (TP).

Focusing on brain dynamics, dynamic FNC (dFNC) analysis shows variations in FNC over time, which give rise to distinct functional integration patterns (dFNC states). The elbow criterion identified four as the optimal number of states. Figure 3 and Figure 4 show the dFNC states. These states are fully reproducible and identified using different clustering procedures (see section 2.32.5). State 1 accounts for 23.76% of all windows (percentage of occurrences, POC = 23.76%), and it is dominated by a strong anticorrelation pattern between the default mode and other functional domains, which can be related to the role of the default mode in reconciling information and subserve the baseline mental activity. State 2 (POC = 38.3%) is distinct by weaker FNC, particularly weaker between functional domains potentially representing the brain’s global segregation state. In contrast, State 3 (POC = 21.31%) demonstrates overall positive FNC across the cerebral cortex, potentially representing global functional integration. Of particular note, the cerebellum shows overall negative FNC with cerebral functional domains in State 3. The negative association between the cerebellar domain and sensorimotor functional domains is prominent in state 4 with POC =16.60%. State 4 can be distinguished with strong functional integration between the visual, somatomotor, and temporal domains, and their anticorrelation patterns with the rest of the brain. This state also shows strong functional integration between the subcortical and cerebellar domains.

### 3.3. Sex-Specific Differences in Individuals with Schizophrenia

Multiscale functional integration was further studied by evaluating sex-specific differences in multiscale sFNC and dFNC between the control group (CT) and the individuals with schizophrenia (SZ). In Figure 3 and Figure 4, columns *J* and *K* show the statistical analysis for each sex cohort using a general linear model (GLM) with age, data acquisition site, and mean framewise displacement as covariates.

In general, sFNC shows more differences between SZ and CT in both sex cohorts than each dFNC state individually; however, the total number of tests that survived FDR correction is comparable between sFNC and dFNC (Supplementary 2). In the female cohort, 576 FNC pairs show significant differences in both sFNC and dFNC, while we identified 638 and 402 FNC pairs show significant differences only in sFNC and dFNC, respectively. In the male cohort, the number of FNC pairs that show significant differences in both sFNC and dFNC is 1076, and the numbers of FNC pairs that show significant differences only in sFNC and dFNC are 720 and 640, respectively. Furthermore, dFNC analysis shows that in the female (male) cohort, 790 (1246) and 3 (21) FNC pairs, respectively, show significant differences in only one dynamic state and all four dynamic states.

Individuals with schizophrenia show reduced sFNC strength within and between the SM and TP domains in male and female cohorts. Looking at dFNC results, we observed these differences emerge in different states for male and female cohorts, i.e., mainly in State 3 for the male cohort and State 4 for females. We observed the sex-specific differences in the SM and TP domains are more pronounced in dFNC states, particularly in State 4. Individuals with SZ also have weaker sFNC and dFNC within the VS and between VS domain and SM and TP domains. Furthermore, with a few exceptions, we observed an overall sFNC and dFNC increases between the SB and the CR, on the one hand, and the SM, the TP, and the VIS on the other hand. We observed the strongest sex-specific differences in State 4 between the VS and the CR.

The results also show significant differences between male and female cohorts across other functional domains in both sFNC and dFNC. For instance, the sFNC between the CC and DMN significant differences in SZ-related alterations between male and female cohorts.

The sex-specific differences are more prevalent in State 4 than sFNC and other dynamic states (Supplementary 2). The number of FNC pairs that show significant sex differences in both sFNC and dFNC is only nine. The results also suggest the largest sex-specific changes in schizophrenia are mainly observed in the dFNC state 4, and they belong to between model-order FNC (Figure 5). Interestingly, we observed opposite patterns of alterations for male and female cohorts in several significant differences. For instance, Figure 5(R1, D4) shows significant differences in the dFNC state 4 (D4) in both male (C2) and female (C3) cohorts. However, while in the male cohort, the strength of dFNC in state 4 reduced in SZ (*t*-value = −3.32), in the female cohort, the strength of FNC increased in SZ cohort (*t*-value = 3.65) compared to control group.

**Figure 5.**
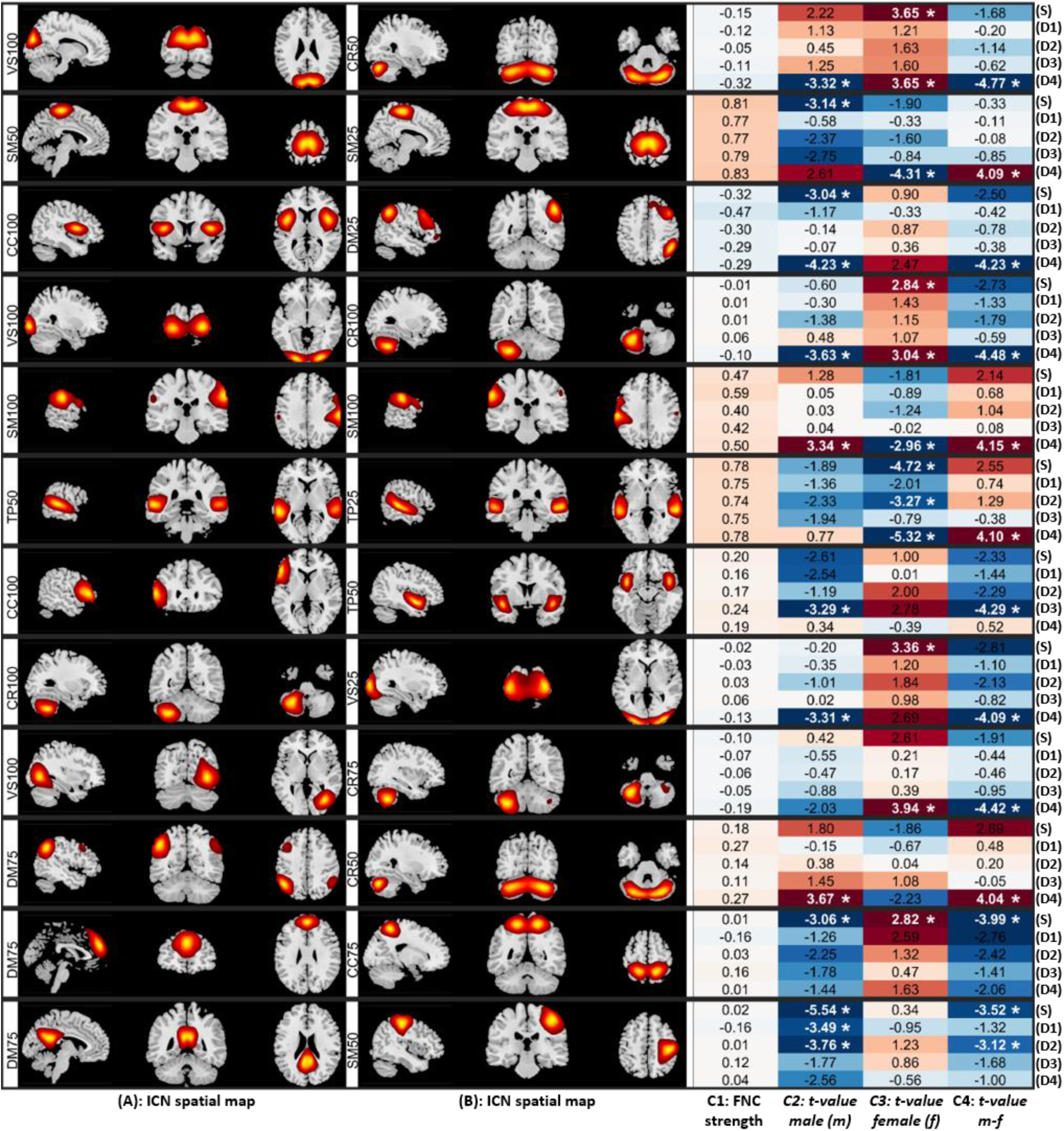
Static and dynamic functional network connectivity (sFNC/dFNC) pairs that show the largest sex-specific multiscale changes in schizophrenia (SZ) presented in twelve rows in order. (S) represents the results of sFNC, and (D1) to (D4) show the results of dFNC for dynamic state 1 to 4, respectively. (A) and (B) display the sagittal, coronal, and axial views of the peak activation of intrinsic connectivity networks (ICNs) associated with each FNC pair. (C1) is the FNC strength. (C2) indicates the t-value of statistical comparisons between typical control and individual with schizophrenia in male cohort. Positive (negative) values indicate stronger (weaker) sFNC/dFNC in individuals with schizophrenia (SZ) compared to the control group. (C3) represents the t-value of statistical comparisons between typical control and individual with schizophrenia in the female cohort, where positive and negative values indicate the same pattern as (C2). (C4) shows the *t*-value of comparing Schizophrenia-related changes between male and female cohorts (“*t*-value of maleSZ vs. maleCT” - “*t*-value of femaleSZ vs. femaleCT”). Asterisk sign * indicates the statistical comparisons that survived multiple comparisons (5% false discovery rate, FDR). Cognitive Control (CC), Default Mode (DM), Visual (VS), Subcortical (SB), Cerebellum (CR), Somatomotor (SM), and Temporal (TP). The number after functional domain abbreviation is the model number; for example, DM25 means the default model domain from ICA model order 25.

One of the advantages of using msICA is that it allows us to see how the same region can contribute to different ICNs at different spatial scales and how the functional connectivity between these ICNs varies across different populations (Figure 5(R2)).

Investigating sex-specific differences at the domain-level across different spatial scales, we observed sex-specific differences are more prominent in the dFNC compared to sFNC. Significant differences exist within the subcortical domain between model order 75 and 100 (SB75-SB100) in sFNC and dFNC state 1 (Figure 6). State 2 shows sex-specific differences between the subcortical and temporal domains within and between several model-orders (Figure 6). State 3, on the other hand, shows sex-specific differences between the cerebellar and somatomotor across different model orders (Figure 6). Like ICN-level comparison, dynamic state 4 reveals the most sex-specific differences, including the temporal, visual, and default mode domains.

**Figure 6.**
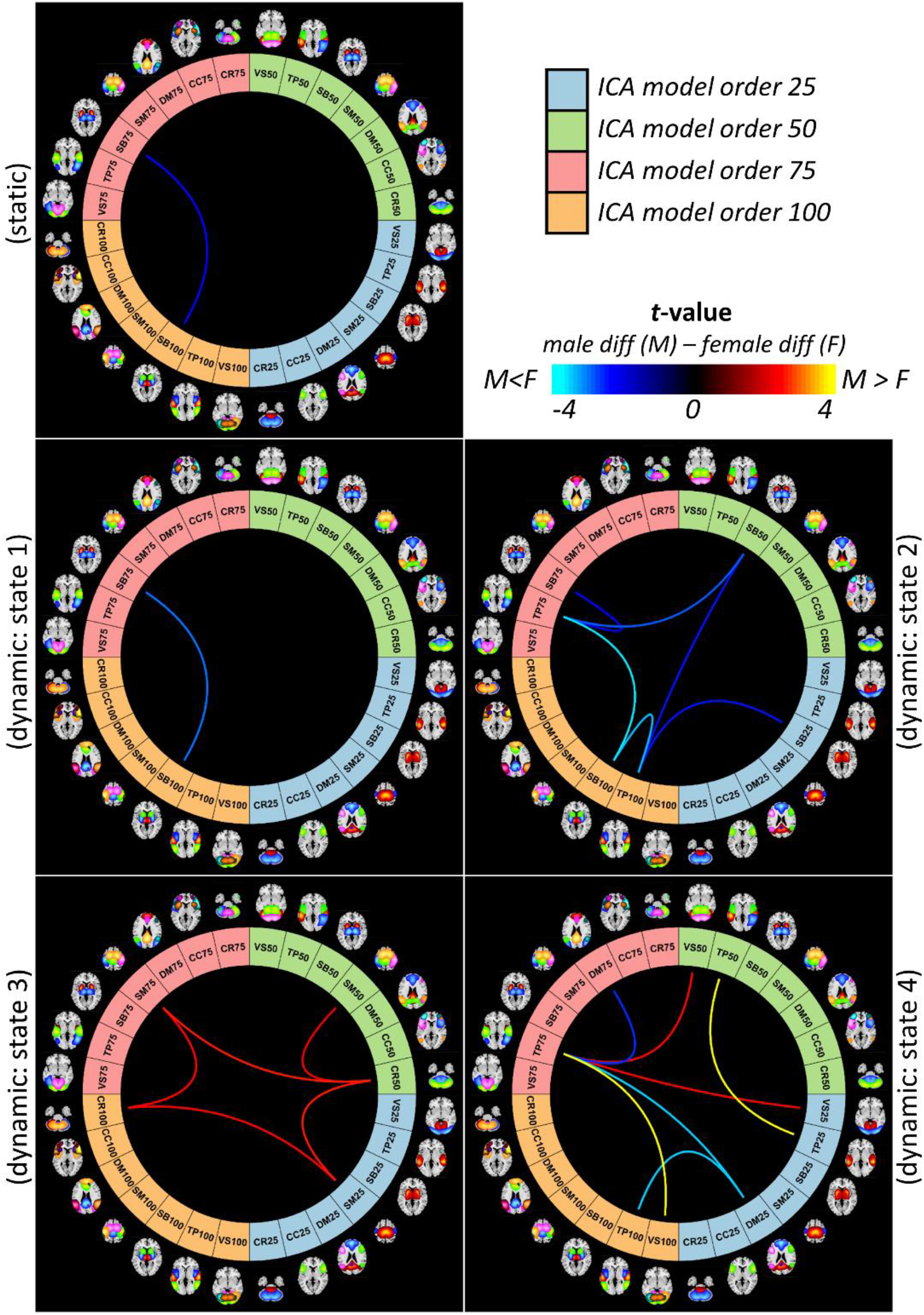
Sex-specific differences at the domain-level across different spatial scales. Cognitive Control (CC), Default Mode (DM), Visual (VS), Subcortical (SB), Cerebellum (CR), Somatomotor (SM), and Temporal (TP). The number after functional domain abbreviation is the model number; for example, DM25 means the default model domain from ICA model order 25.

While sex-specific differences show stronger effects of schizophrenia in males (male diff – female diff > 0) for functional domain connectivity associated with the SM and the VIS, we observe the opposite pattern for the rest of the differences. One exception is the within temporal domain functional connectivity between model order 25 and 50 in the dFNC state 4.

For sex-specific changes at the domain level, we also evaluated the correlation with the PANSS total score. We observed strong correlations with p-value < 0.05 in the male but not the female cohorts for four domain-level features. They include (1&2) within the subcortical domain between model order 75 and 100 (SB75-SB100) in sFNC with the correlation values of 0.281/-0.117 (male/female) and dFNC state 1 with the correlation values of 0.173/-0.197 (male/female); (3) within the temporal domain between model order 25 and 50 (TP25-TP50) in dFNC state 4 with the correlation values of 0.277/-0.062 (male/female); and (4) between the visual domain model order 25 and the temporal domain model order 75 (VS25-TP75) in dFNC state 4 with the correlation values of −0.300/−0.024 (male/female). Among these, SB75-SB100 (sFNC) and VS25-TP75 (dFNC state 4) survived multiple comparison corrections (Supplementary 3).

## 4. DISCUSSION

### 4.1. Multiscale Dynamic Interactions: Functional Segregation and Integration

Studying brain functional connectivity has improved our understanding of brain functions and the impact of brain disorders. However, currently, studying functional connectivity overwhelmingly disregards functional connectivity across multiple spatial scales. Existing studies, at best, apply data-driven approaches like ICA to study functional interactions at single model order but overlook the FNC within and between multiple spatial scales, while the majority of them uses fixed anatomical locations of the same size (e.g., a sphere with the same radius), which in addition to disregarding multiple spatial scales interaction, they ignore differences in the spatial distribution of functional sources.

In this work, we present an approach to study multi-spatial scale dynamic functional interactions, i.e., dynamic changes that occur within and among different spatial scales, a topic that has been overlooked by the field. We leveraged the approach to study schizophrenia’s alterations and its sex-specific differences, which has also been understudied as most schizophrenia research only focuses on single spatial scale FC and non-sex specific alterations of schizophrenia.

### 4.2. Multiscale ICA (msICA)

Our results show that multiscale ICA (msICA) using the Infomax algorithm is an effective, adaptive tool to identify functional sources at multiple spatial scales. Higher model order ICAs segregate the brain and functional domains into more ICNs with, in general, higher spatial granularity. For instance, the subcortical domain splits into more ICNs as the model order increases from 25 to 100. However, ICA does not enforce a limitation on each ICN’s spatial extent. Instead, ICA considers the multivariate association in the BOLD signal to segment the brain. As a result, msICA enables us to visualize functional segregation occurring at different levels of granularity across the brain. This is a desirable characteristic, as we know functional homogeneity varies across the brain and functional domains. The differences in parcellation granularity across functional domains provide additional information about the brain that needs to be studied in the future.

Furthermore, msICA captures the multifunctionality of brain regions and identifies distinct ICNs with high spatial overlap (For example, see the second row of Figure 5). Additional studies are needed to evaluate neurophysiological basis explaining these variations. Furthermore, in this study, we focus on only four model orders of 25, 50, 75, and 100. Future studies should reduce the incremental steps and increase the range of model orders to effectively capture ICNs associated with a larger number of spatial levels of functional hierarchy (Iraji *et al*., 2019d). Recently, we used 1K-ICA, ICA with a model order of 1000, to parcel the brain into very fine-grained functional sources (Iraji *et al*., 2019c). Furthermore, future studies should explore differences across the different back-reconstruction approaches (Erhardt *et al*., 2011). Developing techniques that simultaneously estimate ICNs for multiple model-orders can improve the estimation of ICNs across multiple scales. Finally, considering the recent findings on spatial dynamics (Iraji *et al*., 2019a; Iraji *et al*., 2019d; Iraji *et al*., 2020b), future works should also consider spatial dynamic functional segregations as the spatial patterns of functional sources may vary over time.

### 4.3. Multi-Spatial Scale dFNC

A window-based dFNC approach (Allen *et al*., 2014; Iraji *et al*., 2020a) was adopted to characterize the multi-spatial scale dynamic functional interactions. To our best knowledge, this is the first study that looks at sFNC/dFNC across multiple mode orders. While we observe consistency and similarity of sFNC/dFNC both within- and between-model orders, there are also distinct differences in FNC patterns across FNC patterns. The differences are more distinguishable when there are larger differences in model-orders, e.g., between model orders 25 and 100 (see, for example, Figure 4 (A), (B), and (E)). This further highlights the importance of including a wider range of model-orders in future studies.

Another important point is how we identify dFNC states. In this study, dFNC states were identified using all 127 ICNs; however, the brain may experience different states and/or temporal changes across different spatial scales. Higher functional hierarchy levels have less homogeneity and more dynamic behavior (Iraji *et al*., 2019d). Therefore, we expect more dynamism in the low-model order ICAs. Future work should focus on variation in dFNC states and their timing across multiple model orders and differentiate between global and scale-specific dFNC states. It would also be interesting to extend the same multi-scale idea to the number of clusters for the dFNC analysis. Different cohorts (e.g., male, female, control, and schizophrenia) may depict different characteristics at different scales.

Furthermore, similar to multi-spatial scales, brain functional segregation and integration can occur at different temporal scales and frequencies; thus, future studies can benefit from multi-temporal scale functional interactions. Developing multi-spatiotemporal scale analytic approaches and methodological frameworks to study functional sources is a crucial future avenue of investigation.

Finally, there is a rich repository of dynamic analytical approach and secondary analysis that can be used to evaluate multi-spatial scale brain dynamics (Chang and Glover, 2010; Lindquist *et al*., 2014; Karahanoğlu and Van De Ville, 2015; Yaesoubi *et al*., 2015; Miller *et al*., 2016; Kaboodvand *et al*., 2020).

### 4.4. Schizophrenia

We further investigated the advantage of multi-spatial scale analysis in schizophrenia and identifying sex-specific changes. Our results suggest disruptions in sFNC/dFNC across functional domains. Compared with controls, individuals with schizophrenia show reduced sFNC/dFNC within and between the visual, somatomotor, and temporal domains in both male and female cohorts (Figure 3). Previous studies that looked at differences between typical controls and individuals with schizophrenia also report hypoconnectivity across these functional domains using various approaches (Anticevic *et al*., 2014; Damaraju *et al*., 2014; Kim *et al*., 2014; Shinn *et al*., 2015; Iraji *et al*., 2019a; Iraji *et al*., 2019d; Faghiri *et al*., 2020b). Our study both confirms and extends previous findings. We identify significant differences between males and females in several FNC pairs, mainly showing larger schizophrenia-related changes in males than female cohorts. This can be related to differences in clinical observations, including males presenting more severe overall symptoms, worse outcomes, and slower responses to treatment (Li *et al*., 2016). Greater SZ-related changes across these domains in males are also present at the domain level in dFNC State 4 within the temporal domain and between the temporal and visual domains (Figure 6).

Individuals with schizophrenia show hyperconnectivity of the subcortical domain with the visual, somatomotor, and temporal domains with notable exceptions in dFNC State 4. Unlike sFNC and dFNC in other states, dFNC State 4 has an overall negative association between the subcortical and the visual, somatomotor, and temporal domains (Figure 3). Certainly, temporal lobe anatomical and functional differences have been linked repeatedly to the expression of positive symptoms in schizophrenia (Barta *et al*., 1990; Shenton *et al*., 1992; Woodruff *et al*., 1997). We also observe different patterns of schizophrenia-related changes in male and female cohorts. dFNC State 4 also shows distinct sex-specific differences in the cerebellum domain connectivity patterns, where we observe the opposite pattern of alterations, particularly between the cerebellum and visual domain in the male and female cohorts (Figure 3). Cerebellar dysconnectivity patterns have been linked to negative symptom expression in schizophrenia (Brady *et al*., 2019).

The domain-level analysis suggests that major sex-dependent schizophrenia alterations at a large scale are mainly associated with the subcortical, cerebellar, temporal, and motor domains. Interestingly, most of the sex-specific differences were observed between model-order and associated with dFNC states, highlighting the importance of multiscale dynamic analysis (Figure 4 and Figure 6).

In short, our findings are aligned with and extend previous schizophrenia studies, and we observed explicit sex-specific differences, particularly distinct dFNC patterns in State 4. These demand further investigations into the multi-spatial scale dFNC and sex differences in SZ. However, these findings should be interpreted with caution and considering the limitations of the study.

First and foremost, considering the sex differences in the age of onset, future longitudinal studies should be used to study the role of the age of onset on the sex-specific differences in schizophrenia and evaluate the relationship between time and sex-specific differences over time. Long-term effects of medication and treatment, which cannot be accounted for (Moncrieff and Leo, 2010), might impact observed differences. Including unaffected close relatives sharing genetic risk, i.e., at-high-risk unmedicated subjects, can help us better understand changes in brain function (Pearlson and Stevens, 2020). The unbalanced number of samples between groups is another limitation of the studies. While we control for sex-difference and the null distribution was created with the same female to male ratio, future studies should focus on datasets with larger numbers of females.

### 4.5. Biomarker and Importance of Sex-Specific Characteristics

According to NIH Biomarkers Definitions Working Group, a biomarker is defined as “a characteristic that is objectively measured and evaluated as an indicator of normal biological processes, pathogenic processes, or pharmacologic responses to a therapeutic intervention (Biomarkers Definitions Working Group, 2001)”. As such, a diagnostic biomarker is defined as a characteristic or feature capable of detecting or confirming the presence of a (subtype of) disease or condition of interest (Califf, 2018). At the same time, numerous studies have observed sex differences in schizophrenia, including in the age of onset, in experiencing negative and positive symptoms, and in response to treatments (Nawka *et al*., 2013; Li *et al*., 2016; Seeman, 2019). Therefore, the biomarkers for schizophrenia might be somewhat different for males and females.

This study’s premise is that sex influences differences in schizophrenia characteristics, and we introduce a dynamic multi-spatial scale framework to obtain candidates for sex-specific biomarkers from rsfMRI data. We observed significant sex-specific differences across several functional domains, including in subcortical and temporal connectivity patterns, which also significantly correlate with symptom scores in males but not females. Interestingly, the affected functional domains have been frequently reported to be altered in SZ and touted as having potential to serve as identifying biomarkers. Our results suggest that sex-specific functional connectivity changes might be related to schizophrenia symptoms and underlying causes and emphasize the importance of carefully incorporating sex in the development of diagnostic/predictive/monitoring biomarkers. While sex and schizophrenia can be identified straightforwardly, there has been very little work looking at sex and schizophrenia differences across different spatial scales in resting fMRI data. The incorporation of sex as a biological variable within the context of schizophrenia may help shed new light on the neurobiological mechanisms of schizophrenia and in particular. Future studies should leverage these findings and incorporate sex into feature selection and classification algorithms to identify a set of sensitive schizophrenia-related features for use in updating nosological categories and building diagnostic and predictive models.

## 5. CONCLUSION

Brain dynamic functional interaction can occur at different spatial scales, which has been underappreciated. In this work, we propose an approach that uses multiscale ICA and dFNC to study brain function at different spatial scales. This results in a more comprehensive map of functional interactions across the brain. The not only solves the limitation of using fixed anatomical locations but also eliminates the need for model-order selection in ICA analysis. Therefore, we propose multiscale ICA (msICA), and future multi-spatial scale methods should be broadly applied in future studies. Going forward, we can further improve the proposed approach by incorporating explicit spatial dynamics and multi-temporal scale features of functional sources. We leverage the proposed approach to study male/female common and unique aspects of sFNC/dFNC in schizophrenia, which has not been investigated despite previous reports on sex differences on the prevalence, symptoms, and responses to treatment. The majority of sex-specific differences occur in between-model-order and associated with dFNC states, further highlighting our proposed approach’s benefit. Future studies are needed to validate our findings and evaluate the further benefits of multiscale analysis.

